# Evolutionary analysis of genomes of SARS-CoV-2-related bat viruses suggests old roots, constant effective population size, and possible increase of fitness

**DOI:** 10.1101/2022.02.28.482287

**Authors:** Monika Kurpas, Roman Jaksik, Marek Kimmel

**Affiliations:** Faculty of Automatic Control, Electronics and Computer Science, Silesian University of Technology, Akademicka 16, 44-100 Gliwice, Poland; Department of Statistics, Rice University, 6100 Main Street, Houston, TX 77005, USA

**Keywords:** bat, coronavirus, evolutionary analysis, Moran model, BEAST 2, phylogenetics

## Abstract

It is of vital practical interest to understand the co-evolution of bat *β*-coronaviruses with their hosts, since a number of these most likely crossed the species boundaries and infected humans. Complete sequences of 47 consensus genomes are available for bat *β*-coronaviruses related to the SARS-CoV-2 human virus. We carried out several types of evolutionary analyses using these data. First, using the publicly available BEAST 2 software, we generated phylogenetic trees and skyline plots. The roots of the trees, both for the entire sequences and subsequences coding for the E and S proteins as well as the 5’ and 3’ UTR regions, are estimated to be located from several decades to more than a thousand years ago, while the effective population sizes remained largely constant. Motivated by this, we developed a simple estimator of the effective population size in a Moran model with constant population, which, under the model is equal to the expected age of the MRCA measured in generations. Comparisons of these estimates to those produced by BEAST 2 shows qualitative agreement. We also compared the site frequency spectra (SFS) of the bat genomes to those provided by the Moran Tug-of-War model. Comparison does not exclude the possibility that overall fitness of the bat *β*-coronaviruses was increasing over time as a result of directional selection. Stability of interactions of bats and their viruses was considered likely on the basis of specific manner in which bat immunity is tuned, and it seems consistent with our analysis.

## 1 Background

It seems to be a predominant belief that the recent *β*-coronavirus (Severe Acute Respiratory Syndrome Coronavirus 2, SARS-CoV-2), which causes acute respiratory disease COVID-19, is of zoonotic origin [23]. Other hypotheses, including genetic engineering of the virus, are considered controversial in the research community [2]. For previous epidemics such as the Severe Acute Respiratory Syndrome (SARS) and Middle East Respiratory Syndrome (MERS) caused by *β*-coronaviruses, it has been proven that the reservoir of viruses capable of infecting humans is, among other, the population of bats [23]. It is an interesting question, what is the nature of the interaction of the population of the viruses with bat host.

In this study, we used consensus genomic sequences of bat *β*-coronaviruses to infer the pattern of their evolution and to estimate the relevant parameters. The simulation analysis was supported by a phylogenetic analysis carried out using the Bayesian Evolutionary Analysis by Sampling Trees 2 (BEAST 2) [4] software, which allows the creation of a phylogenetic tree, estimation of the age of the most recent common ancestor (MRCA) of the studied sequences and estimation of size of the studied population using the Bayesian Skyline algorithm. We cross-checked the sophisticated and comprehensive BEAST 2 estimates by using the maximum parsimony analysis [11, 12]. We also developed estimates of mutation rate and effective population size under the hypothesis of neutrality and under sampling that provided genomes captured at different dates.

We also proposed to use a version of the Tug-of-War model introduced by McFarland [18], to model the site frequency spectra (SFS) of the population of viruses and, by comparison with the data, to further refine conclusions from phylogenetic analysis.

In addition, we carried out the analyses using the 3’ and 5’ UTR regions which may include regulatory elements and the genes coding for the envelope (E) and spike (S) proteins. This latter is a glycoprotein that binds to the ACE2 receptor present in the cytoplasmic membrane of the host cells and thus allows the virus to penetrate them [22].

## 2 Methods

### 2.1 Multiple Sequence Alignment and sample preparation

The analysis was carried out using 47 nucleotide sequences of SARS genomes isolated from bat hosts, downloaded from the National Center for Biotechnology Information (NCBI) Virus database [1] (full list of sequences can be found in Supplementary Information). Samples were dated from the period 2004-2017.

The sequences were aligned using MUSCLE multiple sequence aligner [9, 8] based on which we assessed the variability at each genomic position relative to the consensus sequence. For each variant we calculated the fraction of sequences in which an alternative allele was observed, and summarized the data for each gene. Gene positions inside the created consensus sequence were determined using local sequence alignment, based on coordinates of SARS genes obtained from the Reference Sequence (RefSeq) database under accession number NC_004718.3. RefSeq database, created by NCBI, is an open access, annotated and curated collection providing single records for each natural biological molecule (DNA, RNA or protein) for major organisms including viruses, bacteria and eukaryotes. Based on aligned samples, we deduced the ancestral sequence; see further on. Mutation frequency study was carried out in the R computer language, using *seqinr, msa* and *Biostrings* libraries and visualized using *Gviz* and *ggplot2*.

Individual parts of the genome (3’UTR, 5’UTR regions; envelope and spike gene coding fragments) were cut from the alignment based on Multiple Sequence Alignment (MSA) annotations (see Supplementary Information). The length of the complete sequence in the alignment is 30821 nucleotides.

### 2.2 BEAST 2

BEAST 2 (Bayesian Evolutionary Analysis Sampling Trees 2) is a cross-platform program for Bayesian phylogenetic analysis of molecular sequences. It uses Markov chain Monte Carlo (MCMC) method to estimate rooted, time-measured phylogenies. BEAST 2 uses MCMC algorithm to average over tree space, so that each tree is weighted proportional to its posterior probability.

BEAST’s graphical user interface, BEAUti (Bayesian Evolutionary Analysis Utility), allows to specify the inference parameters and create .xml files used for calculation. We performed analyses using the following parameters in corresponding BEAUti booktabs:

- Partitions – allowing browsing uploaded nucleotide/amino acid sequences and division into partitions corresponding to reading frames. In our case nucleotide sequences were either not partitioned (3’UTR, 5’UTR regions) or partitioned (gene E and S; whole genome) in accordance with the assumption of higher mutation rates of third nucleotide in the codon (split {1,2} + 3) with an appropriate reading frame. We linked trees and molecular clocks of both partitions, to generate one set of phylogenetic trees. Site models remained independent.
- Tip Dates – allowing appropriate collection date assignment for each sequence.
- Site Model – allowing choosing substitution model for nucleotide/amino acid sequence. We used HKY (Hasegawa, Kishino, and Yano [16]) substitution model with estimated substitution rate and empirical nucleotide frequencies.
- Clock Model – allowing choosing clock model type. In our inference we used the strict clock model.
- Priors – allowing definition of prior distributions and their parameters. For phylogenetic tree estimation we chose the Coalescent Bayesian Skyline algorithm, which allows reconstruction of population size. For the clock rate prior we chose the Log Normal with parameters {-5, 1.25}, based on BEAST 2 documentation,. The remaining parameters were left as default.
- MCMC – allowing to specify the number of Markov Chain Monte Carlo algorithm steps and to define output files. For each simulation we chose the MCMC step count independently, based on sequence length and final quality of the result.

We analyzed the output from BEAST 2 using Tracer (version 1.7.1) [20], which graphically and quantitively summarizes the distributions of continuous parameters and provides diagnostic information. We used Tracer also to obtain data for Bayesian Skyline plots.

Trees created by BEAST were summarized in TreeAnnotator software. The resulting tree was visualized in FigTree software (version 1.4.4) [19].

### 2.3 Ancestral sequence determination using the maximum parsimony method

#### 2.3.1 *Dnapars* method

In order to obtain ancestral sequence based on available samples we have used *Dnapars* (DNA parsimony method) from PHYLIP package (version 3.698) [11].

This program carries out the unrooted maximum parsimony tree-making algorithm, analogous to the Wagner trees [17], on DNA sequences. The method of Fitch [13] was used to count the number of base changes needed for a given tree. In the maximum parsimony method the phylogenetic tree that minimizes the total number of character-state changes is sought.

The assumptions of *Dnapars* method [10] are briefly listed. Each site evolves independently. Different lineages evolve independently. The probability of a base substitution at a given site is small over the lengths of time involved in a branch of the phylogeny. The expected amounts of change in different branches of the phylogeny do not vary by so much that two changes in a high-rate branch are more probable than one change in a low-rate branch. The expected amounts of change do not vary enough among sites that two changes in one site are more probable than one change in another.

‘The input for this method was the MSA described earlier on. All parameters were left unchanged except the parameter 5: “Print sequences at all nodes of tree”. As a result we obtained the sequences at tree nodes and the ancestral sequence.

#### 2.3.2 Felsenstein’s bootstrap *seqboot* method

The most-parsimonious tree often underestimates the evolutionary change that has occurred. In order to make tree estimation more reliable we used the bootstrap method *seqboot* from PHYLIP package. As an input MSA was used. As an output the set of 100 bootstrapped samples was obtained. In the next step it was used in the *DNApars* method in order to obtain tree estimates with the parameter ”Analyze multiple data sets” enabled.

#### 2.3.3 *Consense* method

In the final step we checked the support for specific forks in consensus tree using *Consense* algorithm from PHYLIP package. The analysis was performed using ”Majority Rule extended” parameter.

### 2.4 Moran Tug-of-War process

We used for simulations a version of the Moran Model, inspired by the Tug-of-War model of McFarland [18]. Consider a population of a fixed number *N* of individuals, each of them characterized by a pair of integers *γ_i_* = (*α_i_, β_i_*), corresponding to the numbers of drivers and passengers in its genotype, respectively. This pair determines the fitness *x_i_* of the *i*-th individual by the formula

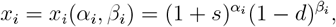

where *s* > 0 and *d* ∈ (0,1) are parameters describing selective advantage of driver mutations over passenger mutations.

This model is a version of the “Tug-of-War” model introduced by McFarland [18]. We provide hypotheses for the complete version. In this paper we will also use the neutral version, with parameters *s* = *d* = 0.

The entire population may be identified with the vector

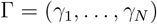

of *N* pairs of integers.

The accompanying vector

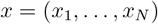

of fitness values determines the future evolution of the population, seen as being under drift and selection pressure, as follows:

The time to the first selection/drift event for the entire Γ is exponentially distributed with parameter (*N* – 1)Σ*x* where

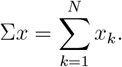

After this time, one individual dies and is replaced by an exact copy of one of the remaining individuals, the probability that the i-th individual dies and is replaced by the *j*-th (*j* ≠ *i*) being 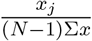. The process then continues with Γ modified by replacing its *i*-th coordinate by its *j*-th coordinate.

Moreover, each individual may, after an independent exponentially distributed time with parameter λ, and independently of other individuals, undergo a mutation event, changing its state to either (*α* + 1, *β*) or (*α, β* + 1) with probabilities *p* ∈ (0,1) and *q* =1 – *p*, respectively.

Mathematical properties of this model are subject of separate publications, in preparation. Here, we used stochastic simulations to explore predictions of this model.

### 2.5 Least square-based fitting of pairwise mismatches in sequence data to the model

One of the purposes of the current analysis was to compare the bat virus sequence data to predictions of a Moran model under neutrality, basics of which are as described in Durrett’s book [7]. We introduced an ad hoc least square method of joint estimation of the effective population size 2*N* and mutation rate in the Moran model with recurrent neutral mutations under the infinite site model (ISM). The method admits observations taken at different times, not only at the bottom of the phylogenetic tree.

The following list (see also Fig. 1) summarizes the notation used.

- *T_i_*, observation time of individual *i* from the sample *i* = 1,2,…,*n*
- *T_lk_* = |*T_l_* – *T_k_*|
- *d_lk_*, count of mutations separating individuals *l* and *k*; under ISM, it is equal to the number of pairwise differences between *l* and *k*
- 2*N*, effective population size, equal to the expected time expressed as generation count, under the Moran model with recurrent mutation and ISM.

**Figure 1:**
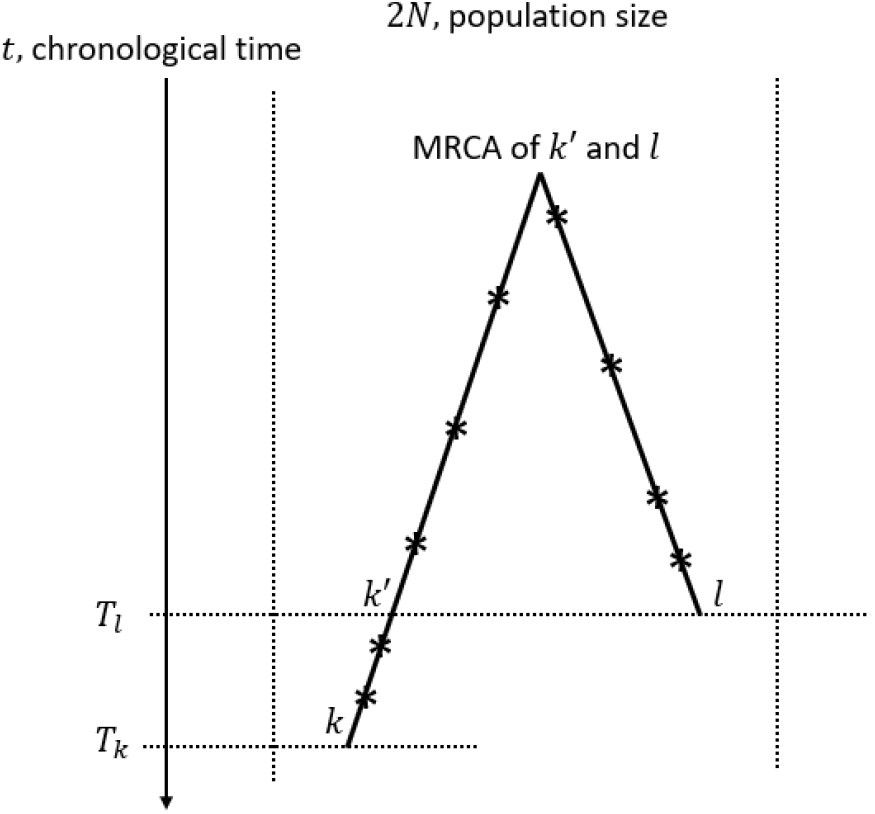
Graphical representation of least-square fitting of pairwise mismatches. Asterisks denote mutation events. Other symbols as in the text.

Expectation of the pairwise difference under assumption that *l* ≠ *k*′ equals to

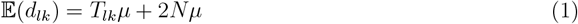

since the expected count of mutations in time *T_lk_* equals *μT_lk_*, while under Moran model and the ISM the expected time to the MRCA equals *N*, so that the expected count of pairwise differences between *k*′ and *l*, equals 2*Nμ*.

Given sample values of *T_k_, k* = 1, 2,…, *n* and *d_lk_*, 1 ≤ *k* < 1 ≤ *n*, we obtain estimating equations for *μ* and 2*N*, by minimizing the sum of squares

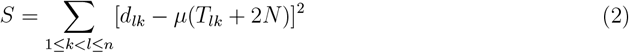

Differentiating with respect to *μ* and 2*N* and equating the derivatives to 0, we further obtain

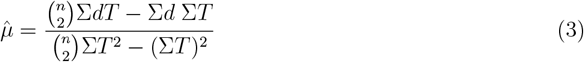

and

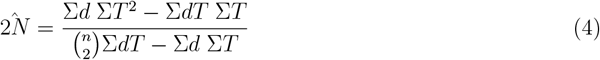

where ∑^*d*^ = ∑_1≤*k<l<n*_ *d_lk_*, ∑*T* = ∑_1≤*k*<*l*≤*n*_ *T_lk_*, *∑dT* = ∑_1≤*k*<*l*≤*n*_ *d_lk_T_lk_*, and 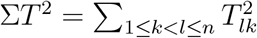

Statistical properties of the resulting estimators were investigated by simulation and proved satisfactory, although there exists a trade-off between estimates of the two parameters (see Appendix).

When applying the method to the selected subunits of the genome we performed sequence quality control to avoid large intervals of deletions or missing nucleotides. We detected such structures in 3’UTR and 5’UTR regions, possibly due to genetic material degeneration or sequencing errors. For this reason we excluded 4 sequences from the 3’UTR and 5 sequences from the 5’UTR dataset. The remaining sequences were trimmed so that they did not contain excess nucleotide deletions.

### 2.6 Site frequency spectrum and its properties

Inference from evolutionary models of DNA often exploits summary statistics of the sequence data, a common one being the so-called Site Frequency Spectrum. In a sequencing experiment with a known number of sequences, we can estimate for each site at which a novel somatic mutation has arisen, the number of genomes that carry that mutation. These numbers are then grouped into sites that have the same number of copies of a mutant. Figure 2 (based on [6]; modified) gives an example with time running down the page. The genealogy of a sample of *n* = 20 cells includes 13 mutational events. We can see that mutations 4, 5, 7, 10, 11, 12, and 13 (a total of 7 mutations) are present in a single genome, mutations 1, 2, and 3 (total of 3 mutations) are present in 3 genomes, mutations 8 and 9 (a total of 2 mutations) are present in six genomes, and mutation 6 is present in 17 genomes. If we denote the number of mutations present in *k* genomes by *S_n_*(*k*), we see that in this example, *S_n_*(1) = 7, *S_n_*(3) = 3, *S_n_*(6) = 2, and *S_n_*(17) = 1, with all other *S_n_*(*j*) equal to 0. The vector (*S_n_*(1), *S_n_*(2),…, *S_n_*(*n* – 1)) is called the (observed) Site Frequency Spectrum, abbreviated to SFS. It is conventional to include only sites that are segregating in the sample, that is, those for which the mutant type and the ancestral type are both present in the sample at that site. Mutations that occur prior to the most recent common ancestor of the sampled genomes will be present in all genomes in the sample; these are not segregating and are called truncal mutations.

**Figure 2:**
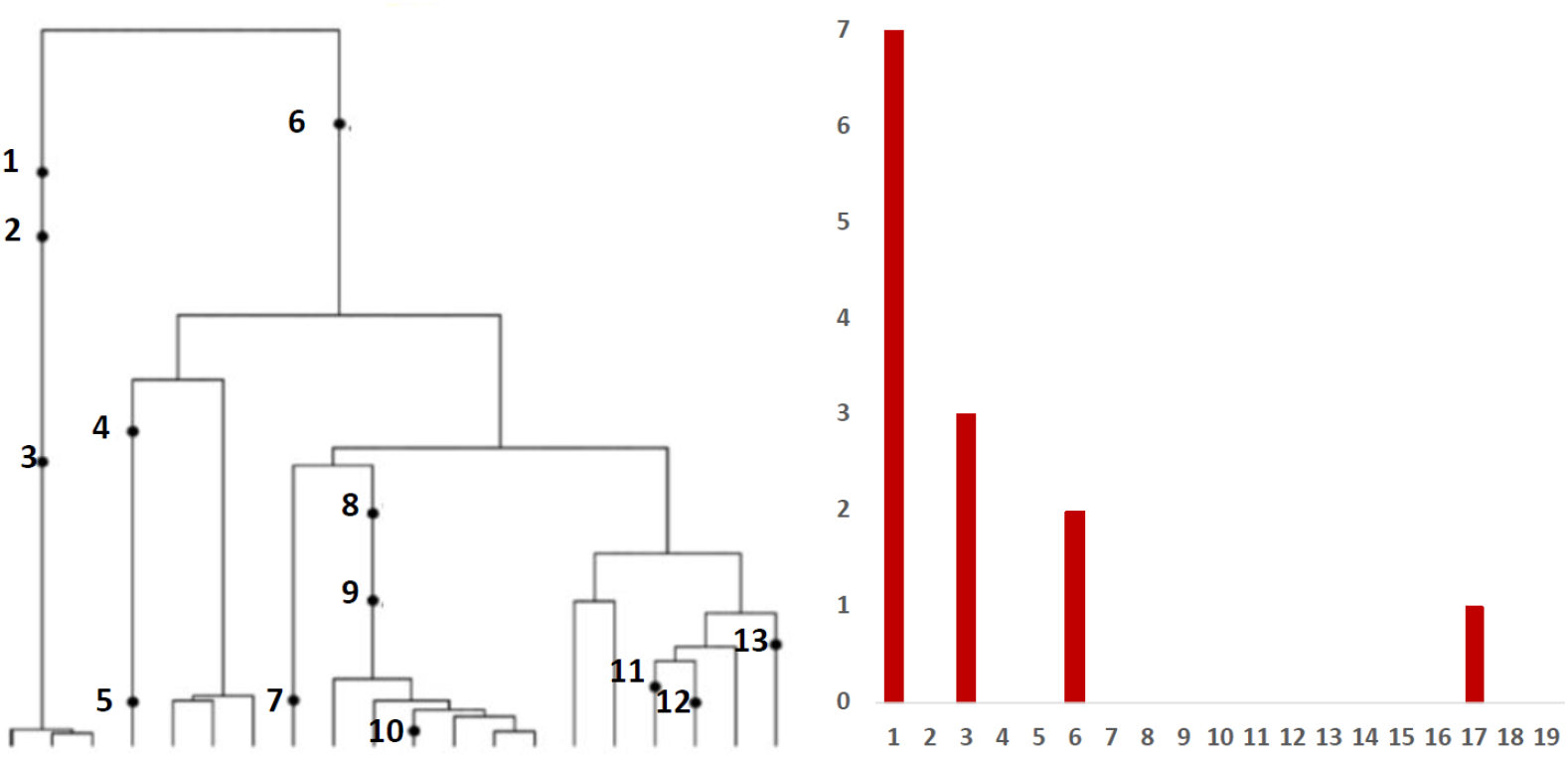
Left panel: Genealogy of a sample of *n* = 20 genomes includes 13 mutational events, denoted by black dots. Mutations 4, 5, 7, 10, 11, 12, and 13 (total of 7 mutations) are present in a single genome, mutations 1, 2, and 3 (total of 3 mutations) are present in three genomes, mutations 8 and 9 (2 mutations) are present in six genomes, and mutation 6 (1 mutation) is present in 17 genomes. Right panel: The observed site frequency spectrum, *S*_20_(1) = 7, *S*_20_(3) = 3, *S*_20_(6) = 2, and *S*_20_(17) = 1, other *S_n_*(*k*) equal to 0.

#### 2.6.1 Site frequency spectrum of bat *β*-coronaviruses

Using MATLAB software we created observed site frequency spectrum of bat *β*-coronaviruses relative to the ancestral sequence obtained with use of PHYLIP package. Except standard nucleotide symbols, the ancestral sequence contained letters that (according to IUPAC rules [5]) represent ambiguity and are used when more than one kind of nucleotide could occur at that position. It also contains “?” symbols, which represents that in addition to one or more bases, a deletion may or may not be present. While evaluating the SFS, we counted mutations in accordance with these rules – if the difference in the sequence at a given site fell within the area of ambiguity we did not count this case as a mutation.

#### 2.6.2 Mathematical properties of site frequency spectra under Moran model with constant population size and neutral infinite site model (ISM)

Griffiths and Tavaré [15] demonstrated that under the Kingman-Wright-Fisher coalescent with constant populations time, the expected SFS of a sample of size *n* from a very large population of size *N* (*n* << *N*), has the form of

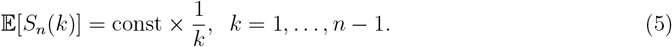

As argued in Dinh et al. [6], the same is the case for the Moral time continuous model, albeit with a different constant. This means that in log-log coordinates the expected SFS under ISM, constant population and neutrality is a straight line with slope equal to −1. As it is known, this is not generally the case when *n* ~ *N*.

## 3 Results

### 3.1 Ancestral sequence determination

We prepared 100 bootstrapped samples using the *seqboot* method from PHYLIP package run on 47 aligned bat *β*-coronavirus sequences. Subsequently we used the *Dnapars* method from the PHYLIP package to create most parsimonious tree for each sample (in some cases more than one tree was found). In order to explore the created set of trees we used the *consense* method from PHYLIP package. The resulting consensus tree (Fig. 3) shows the most probable tree for this dataset. At each node, presented is the number of trees supporting the arrangement of branches displayed, which was determined using the extended majority rule (see Methods).

**Figure 3:**
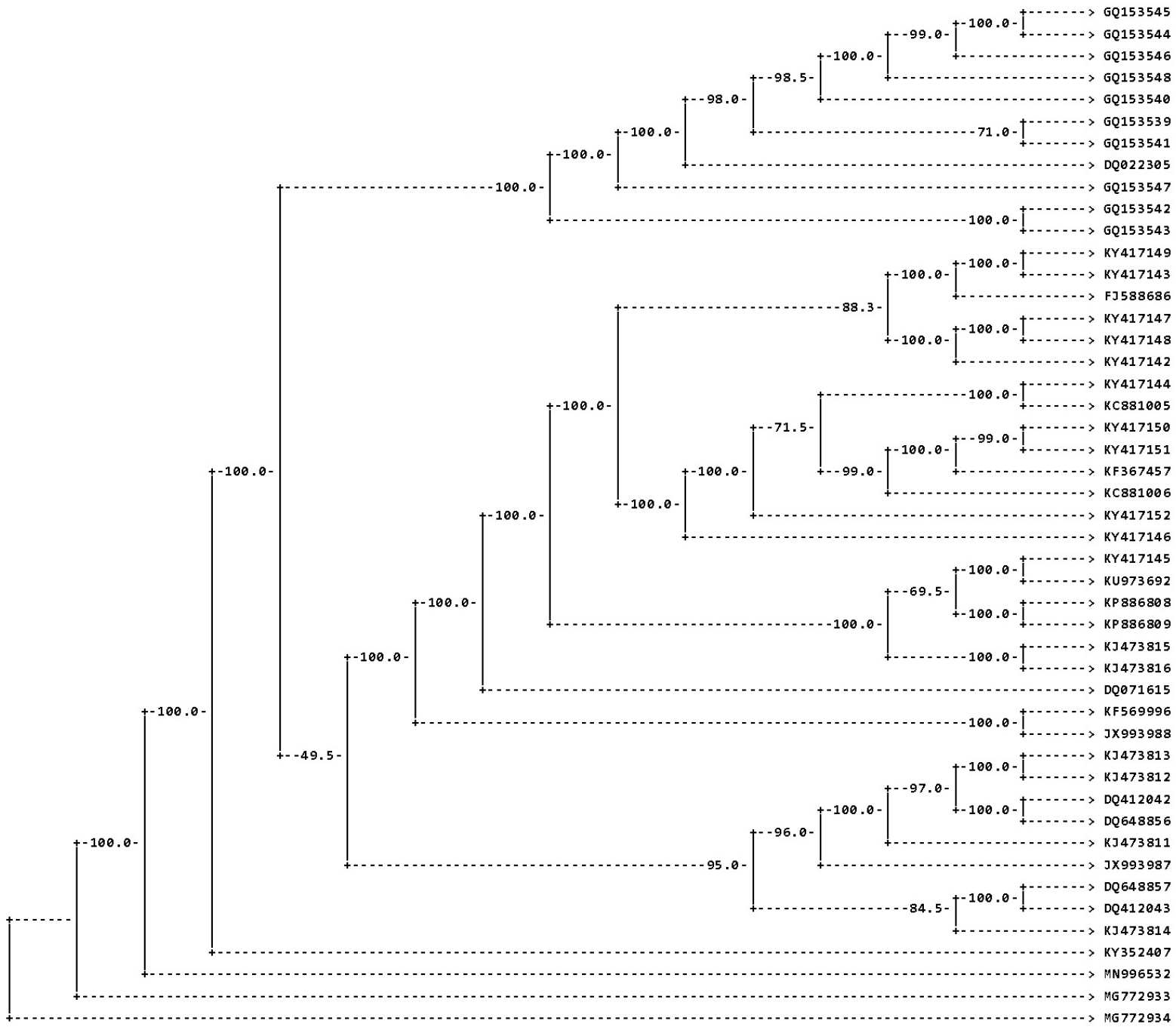
PHYLIP consensus tree for the 47 aligned viral genomes. The leaves are marked with NCBI database identifiers; the numbers on each node correspond to the number of trees having the presented arrangement of branches among 100 datasets created by *seqboot* algorithm.

### 3.2 Genome trees and the most recent common ancestor age

We used BEAST 2 to obtain phylogenetic trees based on the whole genomes (Fig. 4) and four genome fragments (Supplementary Information, Fig. S2, S3, S4, and S5). Based on tip dates applied together with sequences the algorithm is able to estimate the age of most recent common ancestor of aligned sequences. Below we compare estimates for complete sequences (Fig. 5) with sequences trimmed in a way described in section 2.5 (Fig. 6).

**Figure 4:**
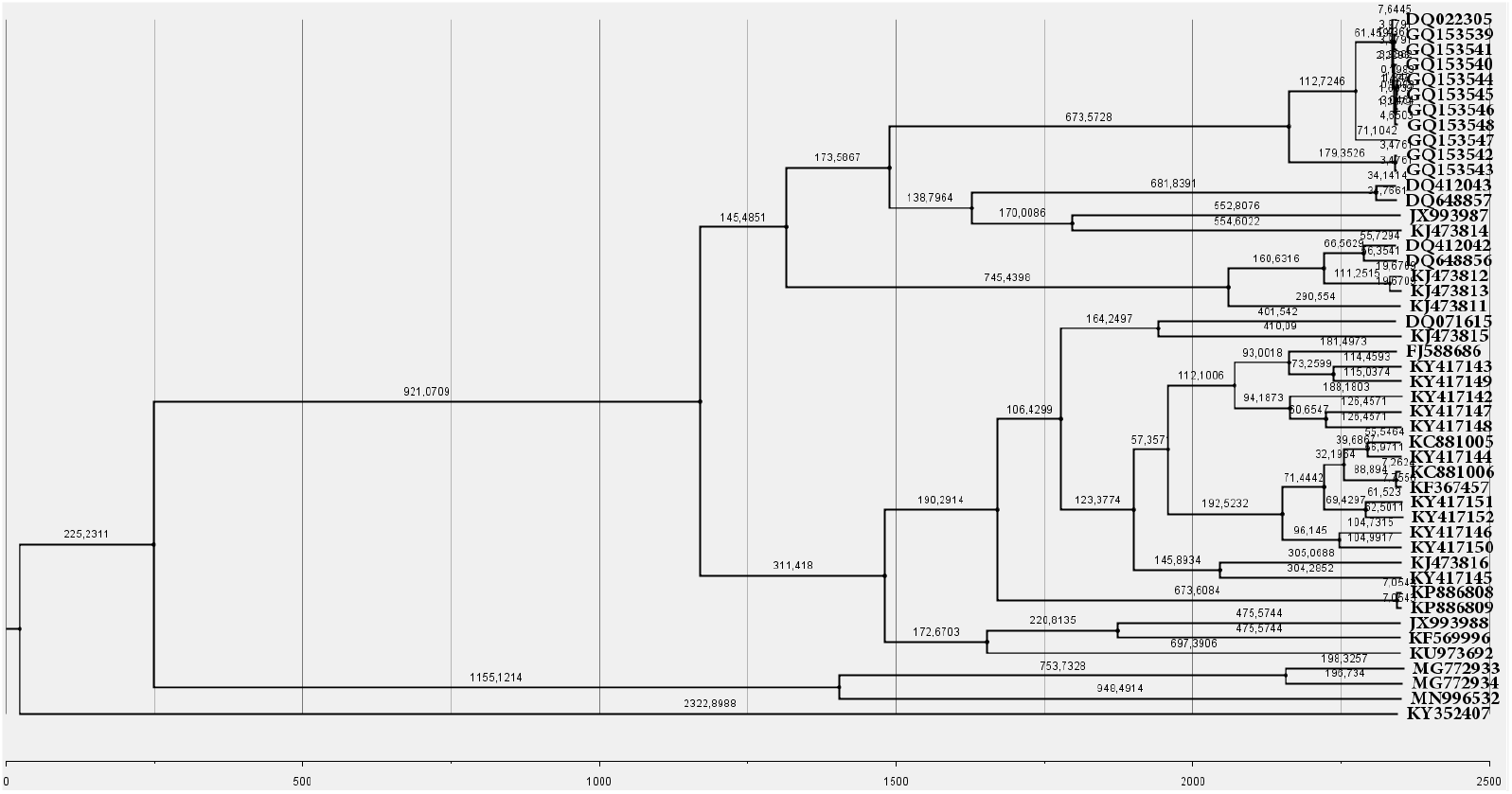
Phylogenetic tree of the whole bat *β*-coronavirus genome inferred by BEAST 2. The leaves are marked with NCBI database identifiers. The numbers on each branch correspond to its length in years. The grid and scale at the bottom of the graph shows the time from the appearance of the MRCA until the date when last sample was sequenced.

**Figure 5:**
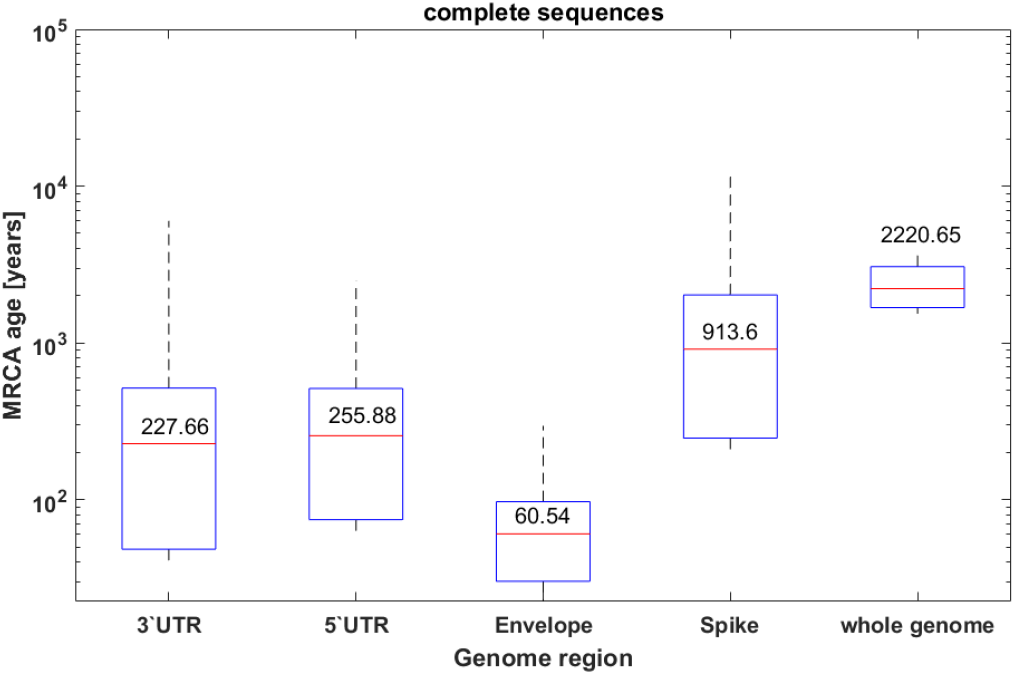
MRCA age (in years) for various parts of genome (complete sequences) inferred by BEAST 2. Boxplots show the mean (red) and 95% highest posterior density (HPD) interval (blue). Whiskers mark the value range.

**Figure 6:**
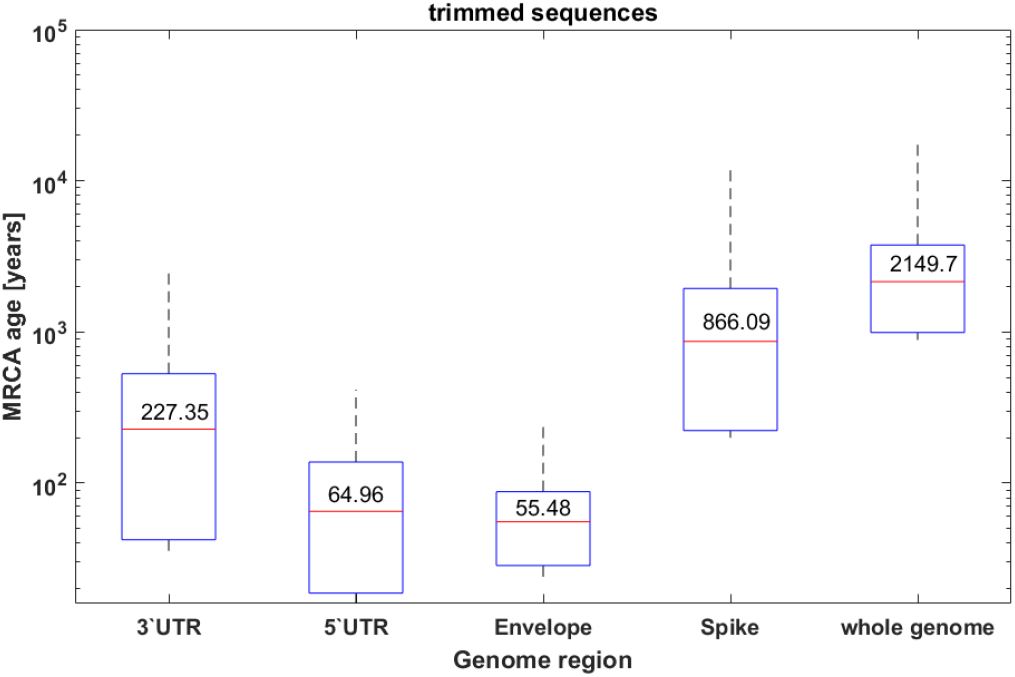
MRCA age (in years) for various parts of genome (trimmed sequences) inferred by BEAST 2. Boxplots show the mean (red) and 95% highest posterior density (HPD) interval (blue). Whiskers mark the value range.

Despite truncation of the sequences, overall MRCA age distribution is similar in both cases and the differences are not statistically significant – boxplots show overlapping interquartile range.

### 3.3 Effective population size and mutation rate estimation

Based on genomic sequences generated as described in Section 2.5 we built the distance matrices and including collection dates for each one of them we calculated estimators using the least square (LS)-based method. Table 1 includes the comparison between MRCA age estimated by BEAST 2 and by our method, which was used also to calculate mutation rate. Let us notice that the estimates of effective population sizes based on the two methods are quite different as are the inferred time scales (see Discussion). However, both methods suggest large age of the most recent common ancestors of the whole genomes as well as the 4 genomic subsets we considered. Also, estimated mutation rates from the least-square method, suggest high conservation of some parts of the genome (the UTR regions and the envelope gene) as opposed to fast-mutating spike gene. Interpretation of these estimates is suggested in the Discussion section.

**Table 1:** Estimates of the MRCA age, effective population size and mutation rate, using the BEAST 2 package, compared to estimates obtained using a direct least-square method.

|  | **BEAST 2:**<br>**MRCA age**<br>[years] | **BEAST 2:**<br>**Population**<br>size from<br>Skyline plots | **LS:**<br>**Population**<br>size = MRCA<br>age in<br>generations | **Generations/year**<br>**= ratio of LS**<br>**to BEAST 2**<br>**MRCA time** | **LS:**<br>**Mutation**<br>rate |
| --- | --- | --- | --- | --- | --- |
| 3'UTR | 227.66 | 200 | 6635 | 29.14 | 0.000569 |
| 5'UTR | 255.88 | 200 | 4687 | 18.32 | 0.000731 |
| E | 60.54 | 40 | 5779 | 95.45 | 0.000858 |
| S | 913.6 | 1400 | 13088 | 14.32 | 0.0203 |
| whole<br>genome | 2220.65 | 2000 | 6550 | 2.95 | 0.3554 |

### 3.4 Population size estimated by Bayesian Skyline

We compared our calculated effective population size with population size estimated by Bayesian Skyline algorithm. In line with our results, BEAST 2 predicted higher effective population size for the whole genome and for the spike gene coding fragment and than for envelope gene or 3’ and 5’ UTR regions (Table 1 and Fig. S6).

In order to check whether the trimming of deletions or missing nucleotides did not affect qualitatively the estimates, we compared Bayesian Skyline plots for complete (Fig. S6 in the Supplement) and trimmed (Fig. S7 in the Supplement) regions of genome. Our results do not show significant changes of shape or estimated values in both cases.

Bayesian Skyline plots in most cases indicated no significant changes in population size over time. We do not observe sharp changes or bottlenecks in the population size.

### 3.5 Moran Tug-of-War model simulations

The Moran Tug-of-War model [18, 7] described in Methods, which includes the stochastic processes of mutation, genetic drift and selection, allows mapping of the evolutionary dynamics occurring in real populations. As explained, some mutations are “driver” mutations that increase fitness of individuals, and other are “passenger” mutations that are neutral or decrease fitness.

In the simulations we first used the neutral version of the model, in which there was no selective advantage of driver mutations over passenger mutations, with parameters *s*=*d*=0 and *p*=0.5, so *v*=*μ*=5, corresponding to average individual lifetime of 0.4 arbitrary time units. Simulated population size *N* was equal to 100 and constant. Total time of simulation *T* was equal to 100 time units (Fig. 7A). We also simulated a number of non-neutral versions of the model, with diverse sets of values of parameters *s* and *d* such as for example *s* = 0.5 and *d* = 0 (Fig. 7B).

**Figure 7:**
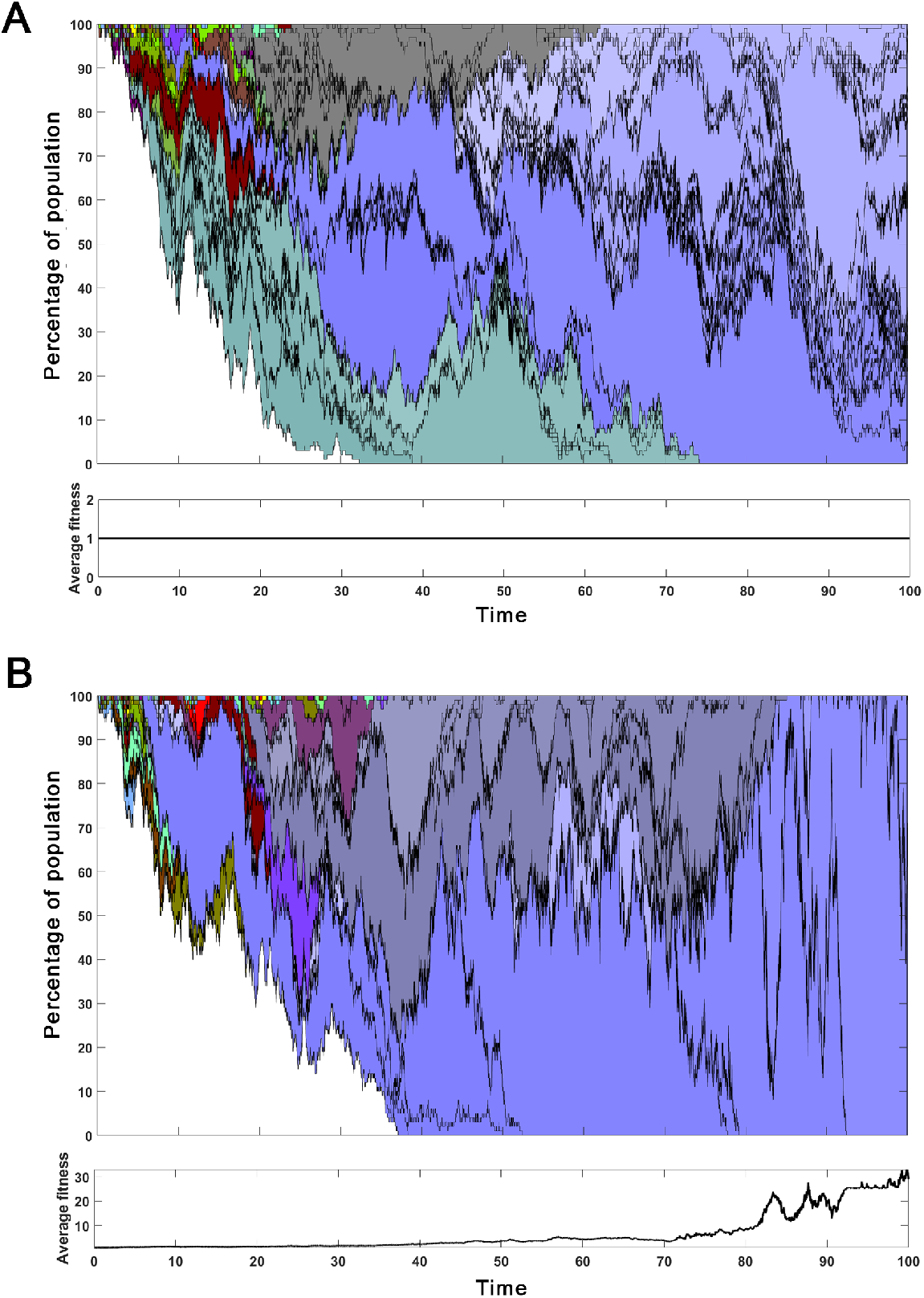
Percentage of given clone in the population of 100 cells acquiring mutations in neutral case (**A**) and in case with selection (**B**). Different shades of the same color distinguish subsequent clones sharing the same parent. Plot on the bottom of each graph shows changes in average fitness per genome over time.

Changes in clonal structure shown in Fig. 7 were visualized using the *area* plot (MATLAB software). At the beginning of the simulation all of *N* =100 cells are the same clone of “zero” type (white color). Direct descendants of clone “zero” are distinguished by new color range. Different shades of a given color correspond to subsequent clones sharing the same parent. Generally speaking, if *ps* > *qd*, the average fitness per genome in the population the population is increasing as evidenced a graph at the bottom of (Fig. 7B) and mathematical considerations (these latter not shown; publication in preparation).

This result show that despite fitness of all clones being equal, we observe significant changes in the clonal structure, with descendants of one (purple) clone taking over the population.

### 3.6 Comparison of observed and simulated site frequency spectra of bat coronavirus genomes

The final analysis performed involves comparison of the SFS of the genomic bat coronavirus spectra to a range of theoretical predictions. As observed in Fig. 8, the empirical spectrum (blue line) is very well approximated by the red dashed line which depicts 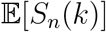 expected from a Moran Model with constant population size under ISM. However, it is almost equally well approximated by the green line, based on the Tug-of-War model with strong positive selection. Other versions of the Tug-of-War model do not fit that well. This apparent contradiction is discussed further on.

**Figure 8:**
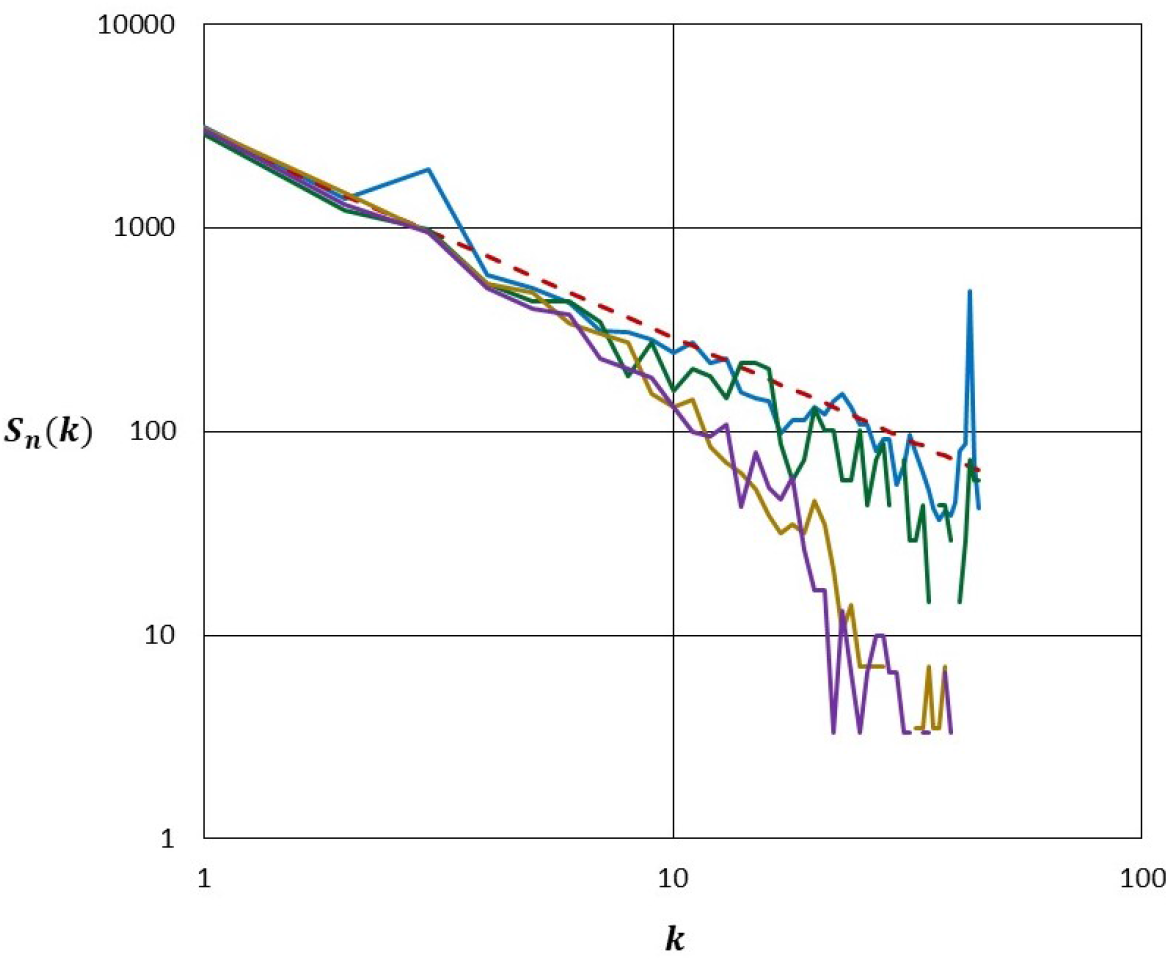
Comparison of site frequency spectra in log-log scale. Red dashed line, 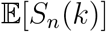 expected from a Moran Model with constant population size under ISM; blue line, SFS calculated from the 47 whole bat coronavirus genomes; green line, Tug-of-War with 100 individuals with parameters *s* = 0.6, *d* = 0, and *p* = 0.5; purple line, Tug-of-War with 100 individuals with parameters *s* = 0, *d* = 0, and *p* = 0.5; pumpkin line, Tug-of-War with 100 individuals with parameters *s* = 0.9, *d* = 0.0001, and *p* = 0.000999.

## 4 Discussion

In this study, we analyzed the genomic sequences of bat coronaviruses, using general use phylogenetic software packages PHYLIP and BEAST 2 as well as the least-square (LS) method and compared these results to predictions of different theoretical models.

As seen in Table 1, the estimated effective population size is smaller for the 3’ UTR and 5’ UTR regions and the envelope gene than for the spike gene, what implicates that the former are more conserved than the latter. This is consistent with the hypothesis that UTRs play a distinctive role in gene regulation, but also with higher mutability with spike, which is seen in the human SARS-CoV-2 genome. This tendency is seen both in BEAST 2 and in the LS estimates. In addition, the Skyline plots (Fig. S6 and S7) suggest approximate time constancy of the bat *β*-coronavirus effective population size. The model underlying the LS estimates directly assumes this constancy.

Differences between effective population size as estimated by BEAST 2 versus the LS and size calculated using proposed method are partly due to the difference in units (years in the case of BEAST 2 and generations the LS method). The definition of generation is unclear in this case, since it does not have to be simply equal to the life expectancy of the pathogen or of the host. One may also interpret the generation time as the time between contacts of infected carriers.

The most striking seem to be the result of the analysis of the site frequency spectrum (SFS) of the 47 genomic sequences. As depicted in Fig. 8, it is quite similar to the classical theoretical prediction [15] under Moran model with constant very large effective population size under ISM (and hence neutrality). However, it seems also consistent with the highly non-neutral version of Tug-of-War model (green line in Fig. 8) with moderate effective population size suggested by BEAST 2 estimates. Since under Tug-of-War with *ps* > *qd*, the overall fitness increases with time, this second possibility might suggest acceleration of evolution of bat coronaviruses.

The present research is only an introduction to the analysis of the co-evolution of *β*-coronavirus genomes and their carriers. If the selective pressure to which viruses are subjected leads to the emergence of new, highly infectious and better adapted strains, then in the long term, it is important to understand and take serious action regarding the potential cyclical recurrences of the current COVID-19 epidemic and subsequent human epidemics.

In bats, the virus-host interaction is regulated by a distinctive action of the innate immune system, different from that in other mammals [3]. Interestingly, the fragments of viral genomes are often permanently inserted into the genomes of their carriers, including the regions undergoing expression and recombination. It is hypothesized that stretches of viral genetic material found in bats’ DNA sequences played a role in the evolution of their immune system [21], which was highly tolerant to pathogens, which may explain why they are considered one of the major reservoirs of viral diseases. As in the organisms of bats, also in humans, the genomes of the virus undergo constant mutations. Research carried out on SARS-CoV-2 sequences has shown that despite the short time that has elapsed since the outbreak of the pandemic, the genetic material contains mutations specific for different regions of the world [14]. Therefore, it is crucial to understand the mechanisms that govern both virus evolution (increasing adaptation to new hosts) and virus-host co-evolution.

## Funding

Monika Kurpas was financially supported by subsidy for the maintenance and development of research potential 02/040/BKM20/1008 (BKM-726/RAU1/2020) granted by Polish Ministry of Science and Higher Education. Roman Jaksik was supported by 2016/23/D/ST7/03665 grant from Polish National Science Centre. Marek Kimmel was supported by the NSF/DMS Rapid Collaborative grant to Marek Kimmel and Simon Tavaré (NSF/DMS-2030577).

## Author’s contribution

MKi suggested the problem, designed and supervised the research. RJ prepared the dataset. RJ and MKu performed the analyses and visualized the results. MKi and MKu prepared the manuscript. All authors reviewed and approved the final version.

## Additional Information

### Competing interests

The authors declare that they have no competing interests.

### Data availability

All relevant data are within the manuscript and its Supporting Information files.

## Supplementary Information

### Accession numbers of 47 genomic sequences of bat SARS-CoV-related *β*-coronaviruses analyzed

KY352407, GQ153542, GQ153543, GQ153547, DQ022305, GQ153548, GQ153546, GQ153544, GQ153545, GQ153541, GQ153539, GQ153540, JX993987, KJ473814, DQ648857, DQ412043, KJ473811, KJ473812, KJ473813, DQ648856, DQ412042, KU973692, KP886808, KP886809, KJ473815, DQ071615, KY417145, KJ473816, KY417151, KY417152, KC881006, KF367457, KY417144, KC881005, KY417146, KY417150, FJ588686, KY417142, KY417143, KY417149, KY417147, KY417148, KF569996, JX993988, MN996532, MG772933, MG772934

### Multiple Sequence Alignment and genome regions annotation

**Figure S1:**
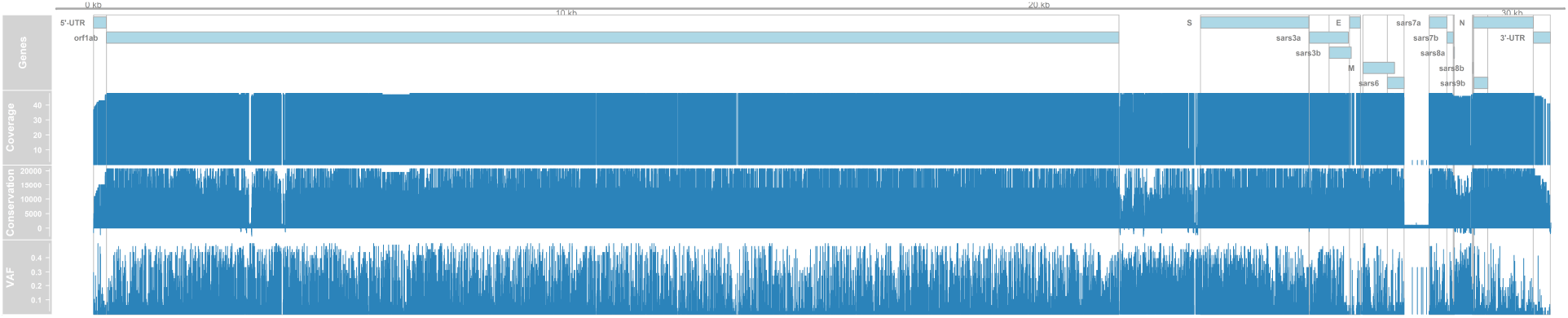
Quality parameters of Multiple Sequence Alignment and genome regions annotation.

### BEAST 2 phylogenies for subsets of genomic sequences

**Figure S2:**
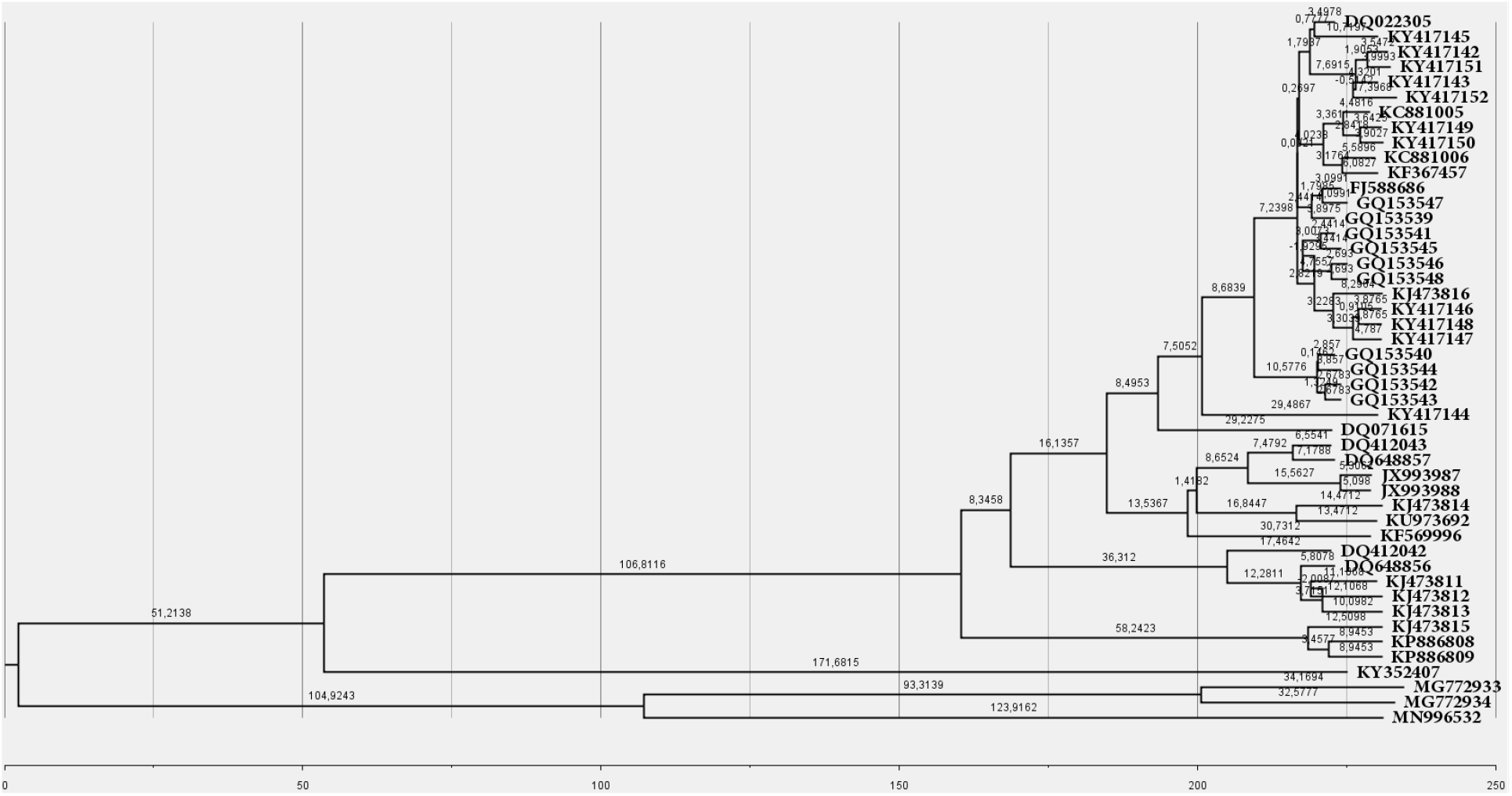
Phylogenetic tree of 3’UTR fragment of bat *β*-coronavirus genome. The leaves are marked with NCBI database identifiers. The numbers on each branch correspond to its length in years. The grid and scale at the bottom of the graph shows the time from the appearance of the MRCA until the date when last sample was sequenced.

**Figure S3:**
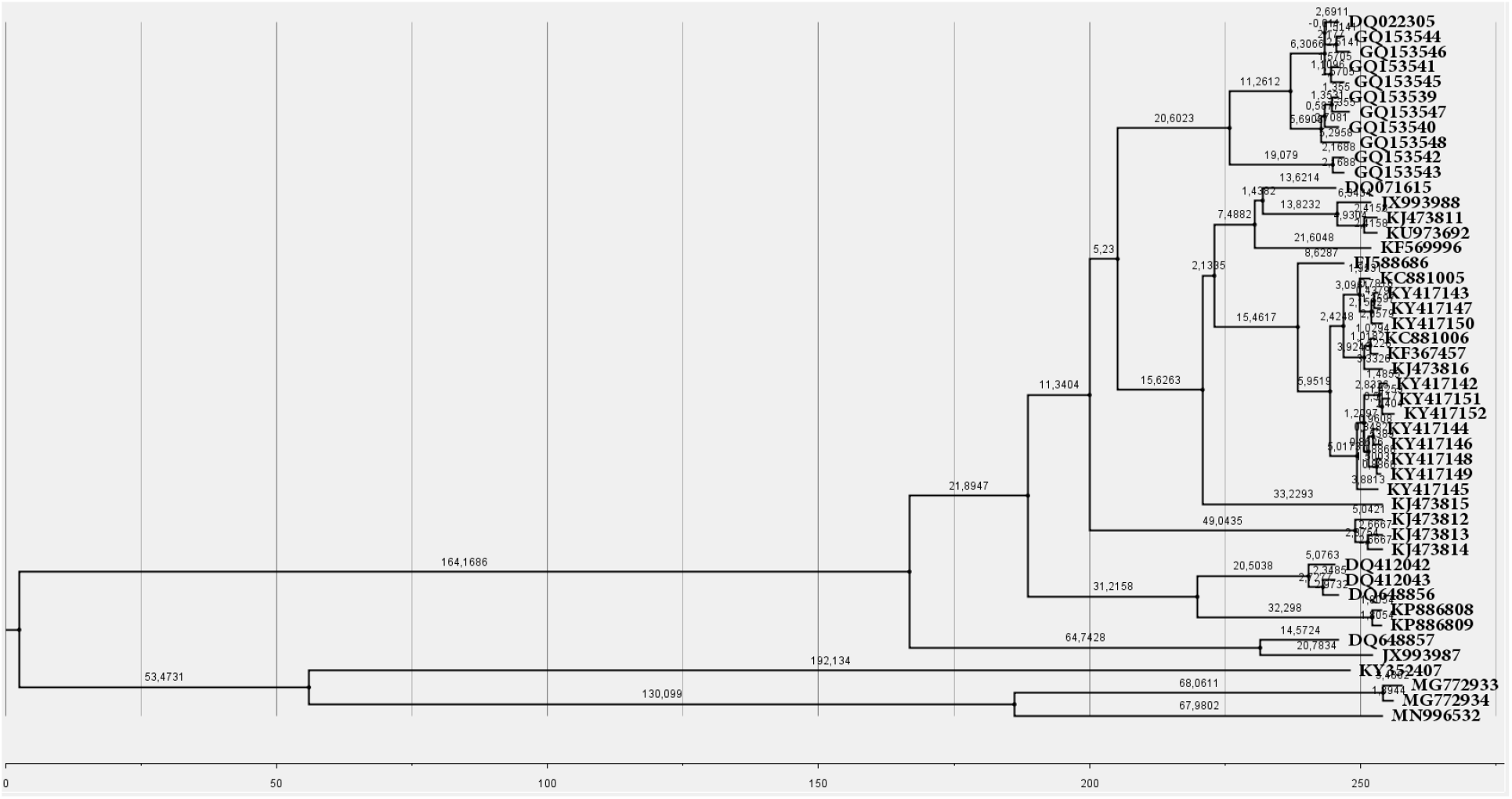
Phylogenetic tree of 5’UTR fragment of bat *β*-coronavirus genome. The leaves are marked with NCBI database identifiers. The numbers on each branch correspond to its length in years. The grid and scale at the bottom of the graph shows the time from the appearance of the MRCA until the date when last sample was sequenced.

**Figure S4:**
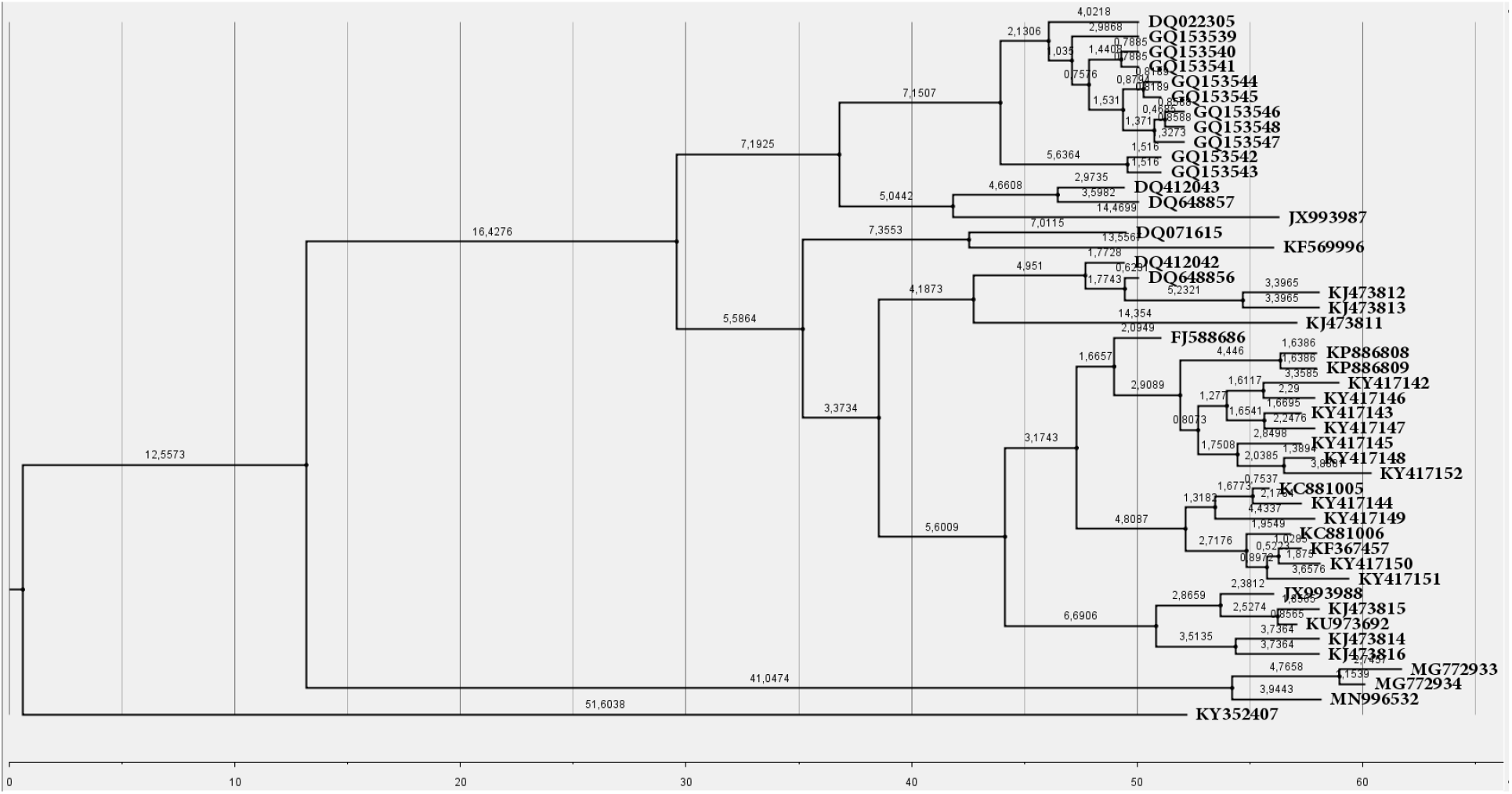
Phylogenetic tree of the fragment encoding envelope gene. The leaves are marked with NCBI database identifiers. The numbers on each branch correspond to its length in years. The grid and scale at the bottom of the graph shows the time from the appearance of the MRCA until the date when last sample was sequenced.

**Figure S5:**
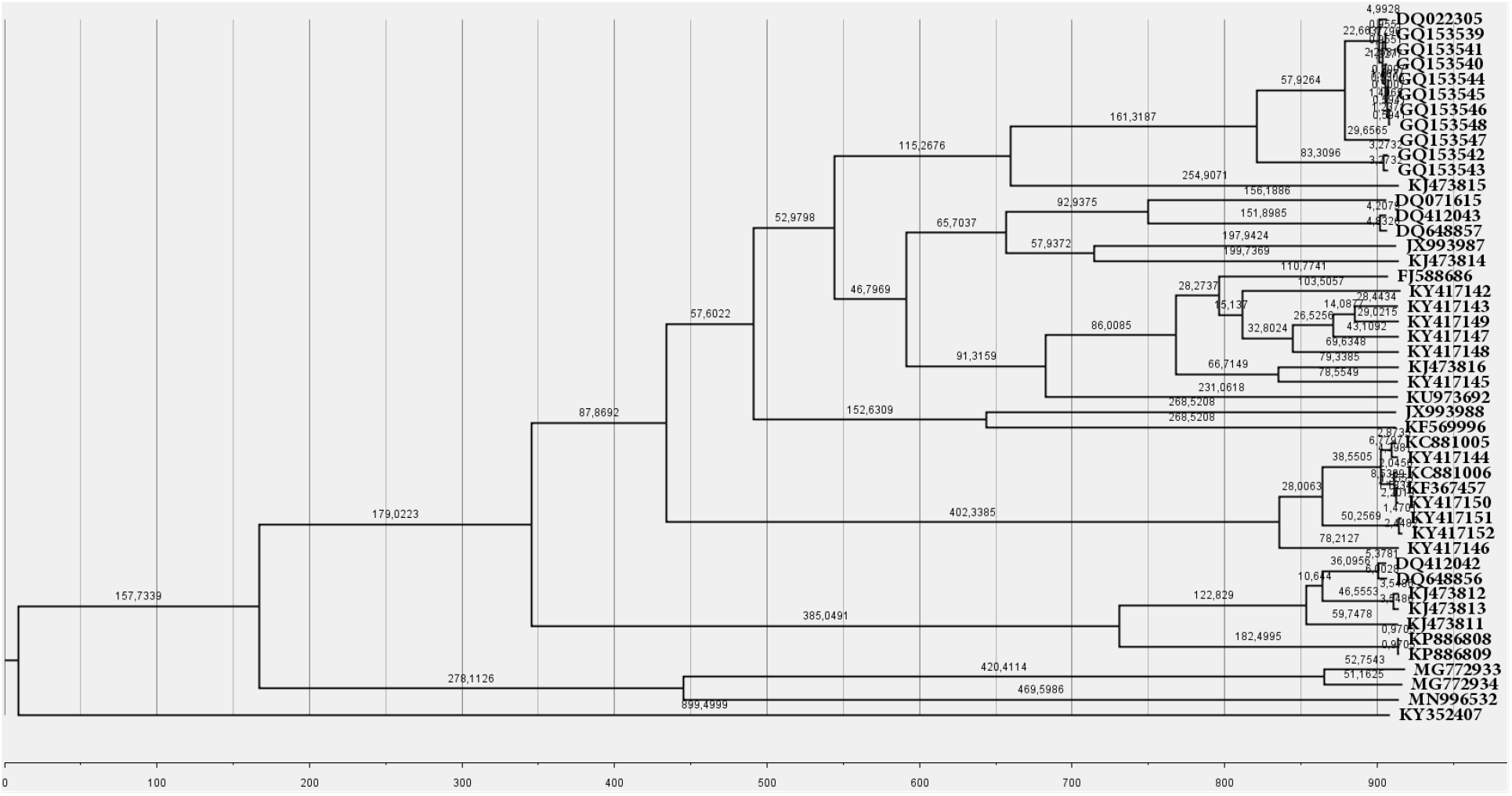
Phylogenetic tree of the fragment encoding spike gene. The leaves are marked with NCBI database identifiers. The numbers on each branch correspond to its length in years. The grid and scale at the bottom of the graph shows the time from the appearance of the MRCA until the date when last sample was sequenced.

### Bayesian Skyline plots

**Figure S6:**
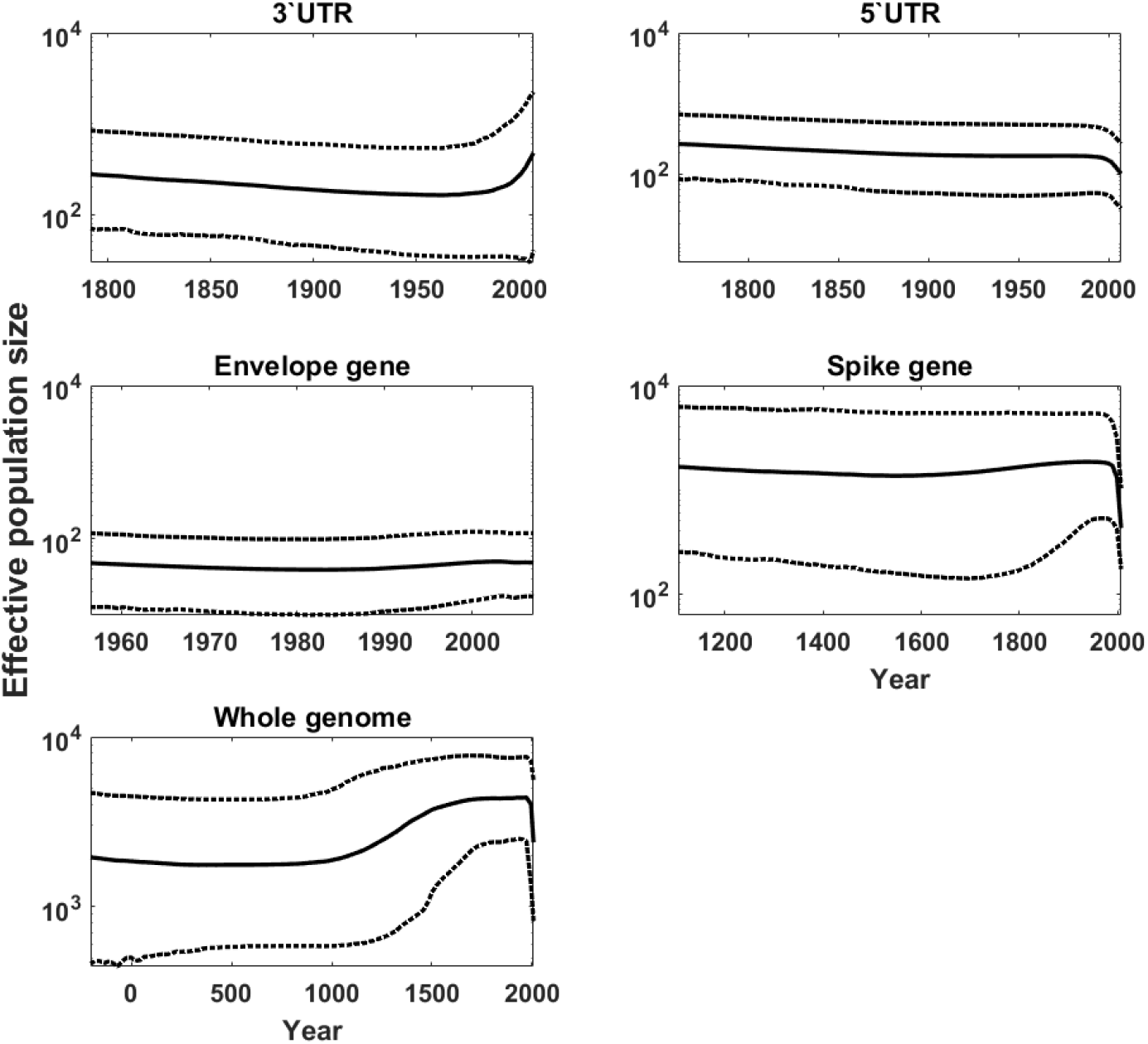
Bayesian Skyline reconstruction of changes in effective population size over time for various parts of the genome (complete sequences). Solid line denotes mean of population size, dotted lines show upper and lower bounds of the 95% HPD interval. The timeline (in years) starts at the MRCA age and ends in 2017 (last collected genome).

**Figure S7:**
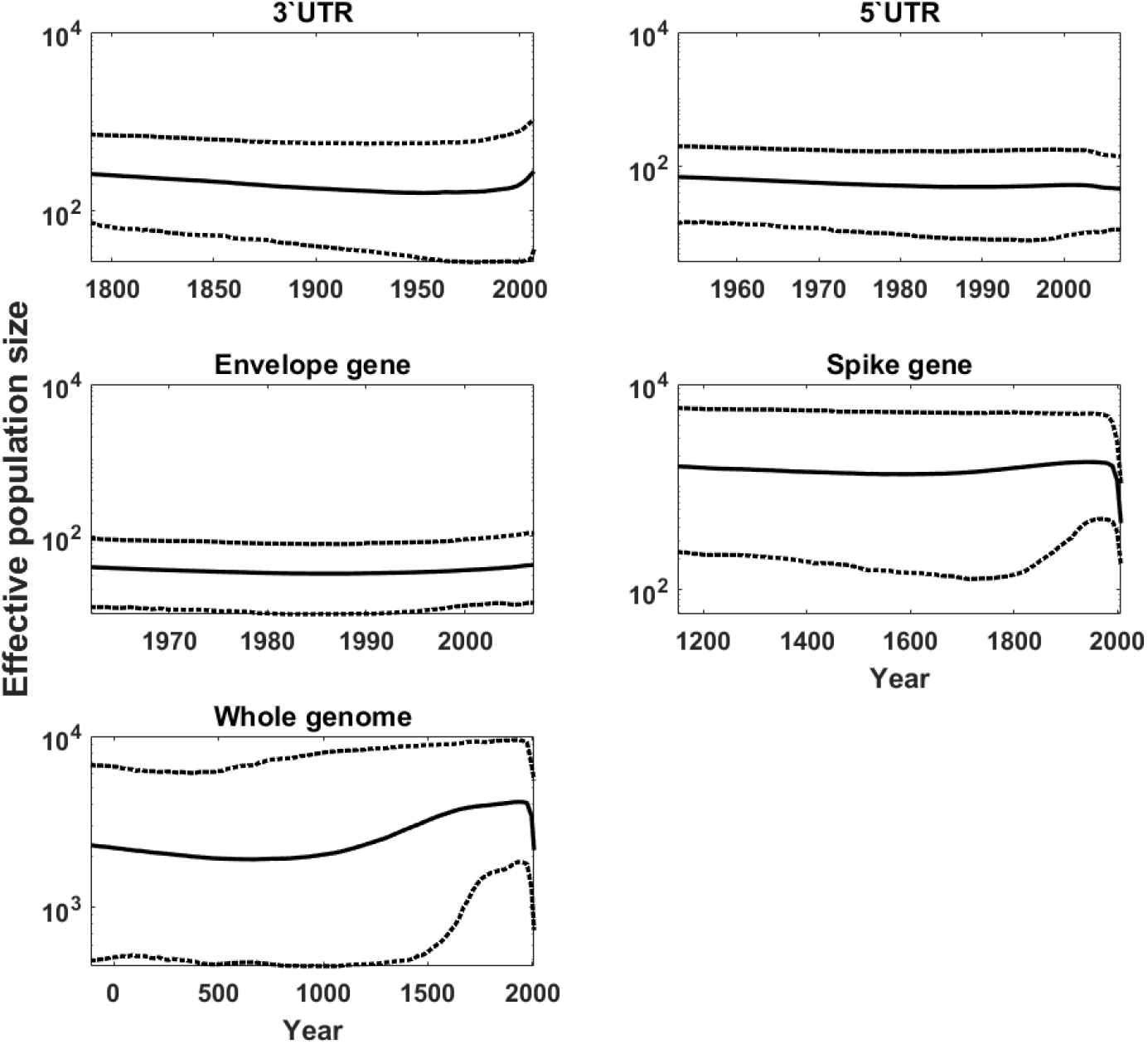
Bayesian Skyline reconstruction of changes in effective population size over time for various parts of the genome (trimmed sequences). Solid line denotes mean of population size, dotted lines show upper and lower bounds of the 95% HPD interval. The timeline (in years) starts at the MRCA age and ends in 2017 (last collected genome).

### Leave-one-fifth-out estimation

We carried out a number of simulations to illustrate the statistical properties of the least-square method. We repeated simulations involving exclusion of random 20% of sequences from the test sample and calculation of the least square estimators based on the remaining ones. These studies show a negative correlation between the estimates of mutation rate and effective population size, this latter equal to the time to MRCA. This effect is expected if the time to MRCA is much higher than the time span of the observations.

The quantity which is best estimable seems to be *θ* = 2*Nμ*. The following graphs (Fig. S8) illustrate results obtained for subsequences of the viral genomes.

**Figure S8:**
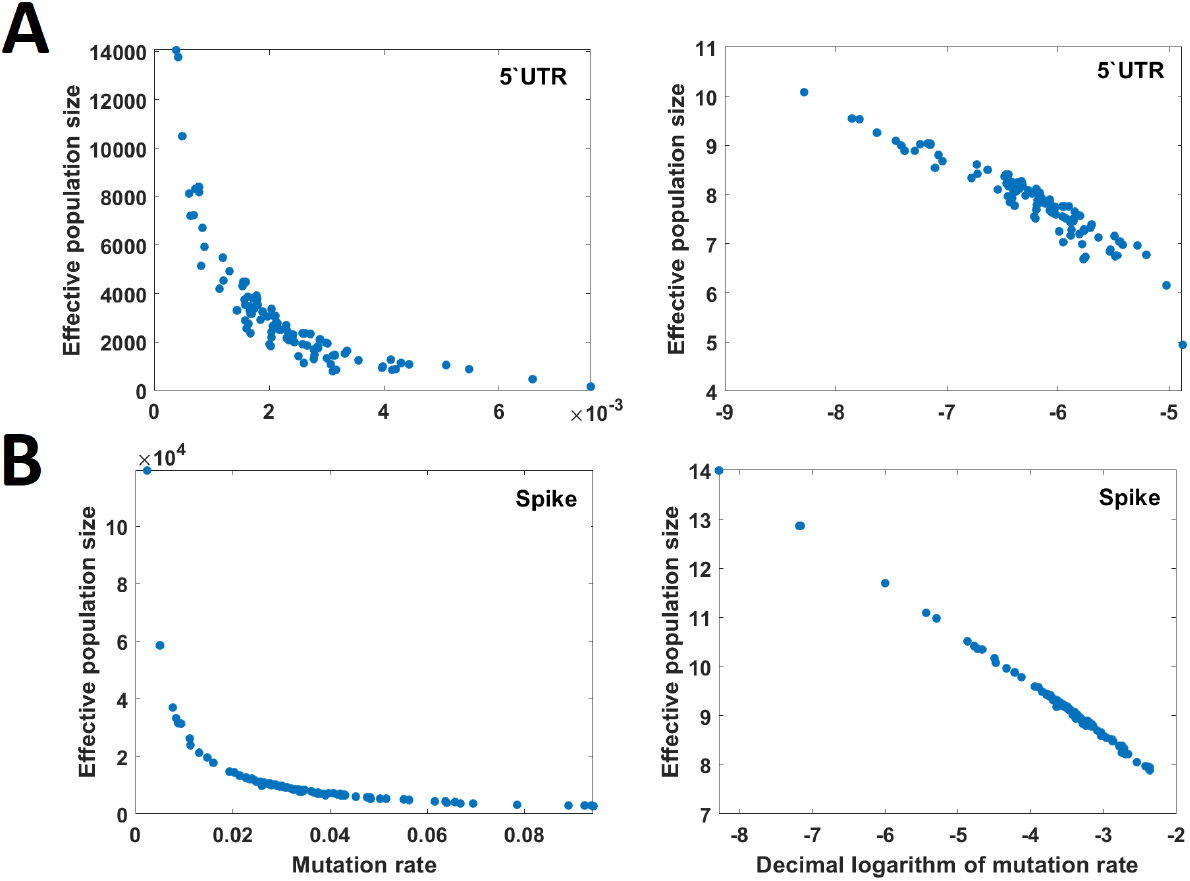
Relationship between effective population size and mutation rate estimates for 5’ UTR region (**A**) and spike gene coding fragment (**B**) in linear (left panel) and logarithmic (right panel) scale. Scatterplots are based on 100 simulations.

**Figure S9:**
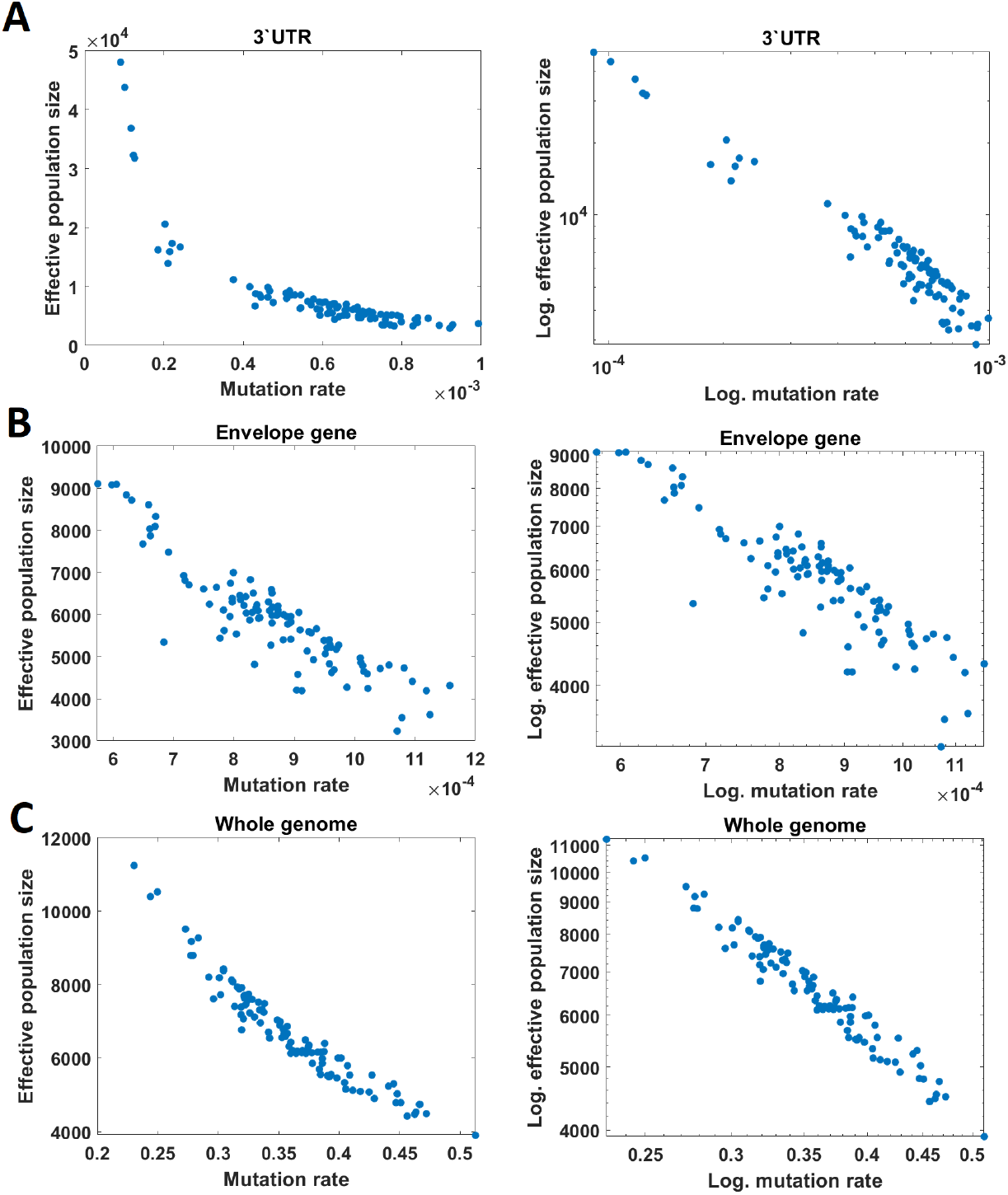
Relationship between effective population size and mutation rate estimates for 3’ UTR region (**A**), envelope gene coding fragment (**B**) and whole genomes (**C**) in linear (left panel) and logarithmic (right panel) scale. Scatterplots are based on 100 simulations.

## Notes

### Competing Interest Statement

The authors have declared no competing interest.

## References

[1] National Center for Biotechnology Information. https://www.ncbi.nlm.nih.gov/. Accessed: 2020-12-30.

[2] Kristian G Andersen, Andrew Rambaut, W Ian Lipkin, Edward C Holmes, and Robert F Garry. The proximal origin of SARS-CoV-2. Nature Medicine, 26(4):450–452, 2020.

[3] Arinjay Banerjee, Michelle L Baker, Kirsten Kulcsar, Vikram Misra, Raina Plowright, and Karen Mossman. Novel insights into immune systems of bats. Frontiers in Immunology, 11:26, 2020.

[4] Remco Bouckaert, Joseph Heled, Denise Kühnert, Tim Vaughan, Chieh-Hsi Wu, Dong Xie, Marc A Suchard, Andrew Rambaut, and Alexei J Drummond. BEAST 2: a software platform for Bayesian evolutionary analysis. PLoS Comput Biol, 10(4):e1003537, 2014.

[5] Athel Cornish-Bowden. Nomenclature for incompletely specified bases in nucleic acid sequences: recommendations 1984. Nucleic Acids Research, 13(9):3021, 1985.

[6] Khanh N Dinh, Roman Jaksik, Marek Kimmel, Amaury Lambert, Simon Tavaré, et al. Statistical inference for the evolutionary history of cancer genomes. Statistical Science, 35(1):129–144, 2020.

[7] Richard Durrett. Probability models for DNA sequence evolution. Springer Science & Business Media, 2008.

[8] Robert C Edgar. MUSCLE: a multiple sequence alignment method with reduced time and space complexity. BMC Bioinformatics, 5(1):1–19, 2004.

[9] Robert C. Edgar. MUSCLE: multiple sequence alignment with high accuracy and high throughput. Nucleic Acids Research, 32(5):1792–1797, 2004.

[10] Joseph Felsenstein. Maximum likelihood and minimum-steps methods for estimating evolutionary trees from data on discrete characters. Systematic Biology, 22(3):240–249, 1973.

[11] Joseph Felsenstein. PHYLIP (phylogeny inference package), version 3.5 c. Joseph Felsenstein., 1993.

[12] Joseph Felsenstein. Inferring Phylogenies, volume 1. Sinauer Associates, Sunderland, MA, 2004.

[13] Walter M Fitch. Toward defining the course of evolution: minimum change for a specific tree topology. Systematic Biology, 20(4):406–416, 1971.

[14] Peter Forster, Lucy Forster, Colin Renfrew, and Michael Forster. Phylogenetic network analysis of SARS-CoV-2 genomes. Proceedings of the National Academy of Sciences, 117(17):9241–9243, 2020.

[15] Robert C Griffiths and Simon Tavaré. The age of a mutation in a general coalescent tree. Stochastic Models, 14(1-2):273–295, 1998.

[16] Masami Hasegawa, Hirohisa Kishino, and Taka-aki Yano. Dating of the human-ape splitting by a molecular clock of mitochondrial DNA. Journal of Molecular Evolution, 22(2):160–174, 1985.

[17] Arnold G Kluge and James S Farris. Quantitative phyletics and the evolution of anurans. Systematic Biology, 18(1):1–32, 1969.

[18] Christopher D McFarland, Leonid A Mirny, and Kirill S Korolev. Tug-of-war between driver and passenger mutations in cancer and other adaptive processes. Proceedings of the National Academy of Sciences, 111(42):15138–15143, 2014.

[19] Andrew Rambaut. Figtree-version 1.4.3, a graphical viewer of phylogenetic trees. Computer program distributed by the author, website: http://tree.bio.ed.ac.uk/software/figtree, 2017.

[20] Andrew Rambaut, Alexei J Drummond, Dong Xie, Guy Baele, and Marc A Suchard. Posterior summarization in Bayesian phylogenetics using Tracer 1.7. Systematic Biology, 67(5):901, 2018.

[21] Emilia C Skirmuntt, Marina Escalera-Zamudio, Emma C Teeling, Adrian Smith, and Aris Katzourakis. The potential role of endogenous viral elements in the evolution of bats as reservoirs for zoonotic viruses. Annual Review of Virology, 7:103–119, 2020.

[22] Renhong Yan, Yuanyuan Zhang, Yaning Li, Lu Xia, Yingying Guo, and Qiang Zhou. Structural basis for the recognition of SARS-CoV-2 by full-length human ACE2. Science, 367(6485):1444–1448, 2020.

[23] Peng Zhou, Xing-Lou Yang, Xian-Guang Wang, Ben Hu, Lei Zhang, Wei Zhang, Hao-Rui Si, Yan Zhu, Bei Li, Chao-Lin Huang, et al. A pneumonia outbreak associated with a new coronavirus of probable bat origin. Nature, 579(7798):270–273, 2020.

